# Changes in perilesional V1 underlie training-induced recovery in cortically-blind patients

**DOI:** 10.1101/2020.02.28.970285

**Authors:** Antoine Barbot, Anasuya Das, Michael D. Melnick, Matthew R. Cavanaugh, Elisha P. Merriam, David J. Heeger, Krystel R. Huxlin

## Abstract

Damage to the primary visual cortex (V1) causes profound, homonymous visual-field loss termed cortical blindness (CB). Though long considered intractable, multiple studies now show that perceptual training can recover visual functions in chronic CB. A popular hypothesis is that training recruits intact extrageniculostriate pathways. Alternatively, training may induce plastic changes within spared regions of the damaged V1. Here, we linked changes in luminance detection sensitivity with retinotopic fMRI activity in eleven chronic CB patients, before and after extensive visual discrimination training. Our results show that the strength of spared V1 activity representing perimetrically blind-field locations before training predicts the amount of training-induced recovery of luminance detection sensitivity. Additionally, training caused an enlargement of population receptive fields in perilesional V1 cortex, which increased blind-field coverage. These findings uncover fundamental changes in perilesional V1 cortex underlying training-induced restoration of conscious luminance detection sensitivity in cortically-blind patients.

---

The primary visual cortex (V1) is the chief cortical relay of visual information from retino-geniculate centers towards higher-level extrastriate areas. Unilateral damage to V1 or its immediate afferents (optic radiation) causes cortically-induced blindness (CB)–a homonymous loss of conscious vision over the contralateral hemifield. Strokes (either ischemic or hemorrhagic) involving the posterior or middle cerebral arteries account for the majority of cases (Zhang et al., 2006a, b). Although CB patients often show damage to extrastriate visual areas, it is damage to V1 that seems to cause the loss of conscious vision and the appearance of defects in luminance contrast detection, routinely measured using clinical perimetry devices (Leopold, 2012; Melnick et al., 2016b). The prevalence of blindness-inducing post-chiasmal lesions of the visual system in the general population is remarkably high (Pollock et al., 2011), and its impact on everyday life deeply debilitating (Papageorgiou et al., 2007). Yet, there is a lack of validated clinical therapies that can help restore, rather than compensate for, deficits in CB patients (Ajina and Kennard, 2012; Chokron et al., 2008; Melnick et al., 2016b; Perez and Chokron, 2014; Pollock et al., 2011; Urbanski et al., 2014). In stark contrast with well-established physical therapies prescribed following motor cortex strokes, the ability of restitution therapies to restore vision in CB patients remains controversial.

Spontaneous recovery typically occurs during the first 3 months post-lesion, with little improvement observed following this phase, and significant changes no longer observed in chronic patients, >6 months post-lesion (Zhang et al., 2006a, b). The first major commercial restoration treatment for chronic CB patients–NovaVision’s Visual Restoration Therapy (VRT)(Kasten et al., 1998)– proposed that intensive computer-based training on a microperimetric luminance detection task could help restore visual sensitivity and shrink perimetrically-defined blind fields in CB patients. Although early findings generated strong interest, subsequent work identified flaws in NovaVision’s approach, which when corrected, revealed it to be ineffective (Reinhard et al., 2005; Schreiber et al., 2006). Despite this initial failure, which reinforced the clinical dogma, scientific teams worldwide subsequently showed that intensive training with gaze-contingent stimulus presentation inside the blind field can restore a range of visual functions at trained, blind-field locations (Cavanaugh et al., 2017; Cavanaugh and Huxlin, 2017; Cavanaugh et al., 2015; Chokron et al., 2008; Das et al., 2014; Herpich et al., 2019; Huxlin et al., 2009; Melnick et al., 2016b; Perez and Chokron, 2014; Raninen et al., 2007; Sahraie et al., 2010; Sahraie et al., 2006). Critically, binocular eyetracking, gaze-contingent stimulus presentation during pre- and post-training tests ensured that improvements within the blind field could not simply be explained by the development of compensatory eye movement strategies. Rehabilitation studies in chronic CB have in common the fact that, unlike perceptual learning in intact hemifields, significant improvements at blind-field locations require daily, retinopically-specific training over weeks to months. This and other differences between training in intact *versus* damaged brains remain a mystery, partly due to our poor understanding of the capacity of damaged, adult visual systems for perceptual processing, plasticity and ultimately learning. Given that we now possess the means to reliably induce recovery in CB, we are in an ideal position to assess the neural mechanisms underlying this phenomenon in V1-damaged individuals.

One proposed recovery mechanism is that training stimulates and improves visual processing in extrageniculostriate pathways mediating *blindsight* (Ajina and Kennard, 2012; Chokron et al., 2008; Das and Huxlin, 2010; Perez and Chokron, 2014; Smirnakis, 2016). Both human and non-human primates with V1 lesions exhibit visually-guided perceptual abilities within their blind field, despite lacking awareness (Ajina and Bridge, 2016; Leopold, 2012; Weiskrantz et al., 1974). Because blindsight is elicited by large stimuli with high-temporal and low-spatial frequency content, it is thought to rely mainly on direct geniculo-hMT+ and superior colliculus-pulvinar-extrastriate projections (Ajina and Bridge, 2016; Ajina and Kennard, 2012; Ajina et al., 2015; Bridge et al., 2008; Leopold, 2012; Schmid et al., 2010; Weiskrantz et al., 1974). Accordingly, some rehabilitation approaches have targeted blindsight to help CB patients recruit these unconscious processes, with some evidence of increased visual performance and awareness post-training (Chokron et al., 2008; Melnick et al., 2016b; Perez and Chokron, 2014; Sahraie et al., 2010; Sahraie et al., 2006). However, training can also recover the ability to discriminate visual information with spatio-temporal properties and motion integration requirements that fail to elicit blindsight (Cavanaugh et al., 2017; Cavanaugh and Huxlin, 2017; Cavanaugh et al., 2015; Das and Huxlin, 2010; Das et al., 2014; Huxlin et al., 2009; Melnick et al., 2016b; Saionz et al., 2019). Thus, “blindsight pathways” may not be the only means by which training can elicit recovery. This is particularly encouraging given the restricted bandwidth of blindsight abilities and the fact that not all CB patients exhibit blindsight.

An alternative hypothesis is that recovery in CB relies on spared, perilesional V1 regions that are functionally impaired, but can be brought back “online” using visual training (Cavanaugh and Huxlin, 2017; Das and Huxlin, 2010; Huxlin et al., 2009; Papanikolaou et al., 2014; Smirnakis, 2016). Here, we directly tested a potential role of spared V1 cortex in training-induced recovery of luminance sensitivity in eleven adult patients with chronic, stroke-induced CB. To do so, we compared retinotopic fMRI activity in V1 with changes in luminance detection sensitivity measured with Humphrey Visual Field (HVF) perimetry, before and after training. By relying on stringent inclusion criteria and a more uniform patient group than most prior studies, we identified ubiquitous, functional changes mediating recovery in chronic, stroke-induced CB. We show that training-induced recovery of luminance detection sensitivity in chronic CB relies upon changes in perilesional V1 regions with spared, visually-evoked, blind-field activity prior to training. As such, our results provide vital insights regarding what may be ubiquitous neural mechanisms mediating visual restoration following long-standing V1 damage.

## Results

All eleven CB patients in the present study (see **Table 1** for patient demographics) had long-standing (35±68 months, range: 5-237 months) unilateral, homonymous, hemianopic visual-field defects secondary to unilateral strokes affecting the occipital cortex. Stroke-induced damages to the posterior cerebral artery and its territory resulted in lesions primarily located in the medial aspect of the occipital lobe and calcarine sulcus (average lesion volume: 16,673±18,592 mm^3^, range: 667-64,320 mm^3^). The precise extent of the lesions into the cuneus and the lingual gyrus varied across patients, as did the degree of ventricular enlargement, suggesting differential involvement of the optic radiations and occipital white matter tracts. Anatomically, the occipital pole was intact in all cases, consistent with preserved foveal sensitivity (35.7±0.85 dB HVF sensitivity) and ability to fixate precisely (measured with eye-tracking).

**Table 1.** Patient demographics. All 11 patients suffered from unilateral, stroke-induced damage to the occipital cortex at least 5 months before training onset, causing a loss in luminance detection sensitivity assessable using Humphrey’s Visual Field (HVF) mapping. Pre-training percent deficit size is computed relative to either a simulated, full hemifield loss or for the inner ±11.5deg area stimulated during fMRI in the present study. Following visual discrimination training (static orientation, motion direction, or both), all patients showed HVF improvements.

| Patient | Gender | Age (years) | Damaged hemisphere | Lesion (mm<sup>3</sup>) | Time post-lesion (months) | Pre-post interval (months) | Pre-training deficit size (deg<sup>2</sup>) |  | Post-training improvement (deg<sup>2</sup>) |  | Training regime |
| --- | --- | --- | --- | --- | --- | --- | --- | --- | --- | --- | --- |
| | | | | | | | full HVF | inner $\pm 11.5$ | full HVF | inner $\pm 11.5^\circ$ | |
| CB1 | M | 67 | L | 6,422 | 10.4 | 5.5 | 692.5 (86%) | 133.4 (51%) | 121.9 | 40.18 | double |
| CB2 | F | 58 | R | 16,303 | 11.8 | 10.8 | 293.6 (37%) | 115.5 (44%) | 317.8 | 85.28 | static |
| CB3 | M | 63 | L | 667 | 38.4 | 34.4 | 472.7 (59%) | 99.9 (38%) | 65.5 | 51 | double |
| CB4 | F | 29 | R | 2,126 | 5 | 3.1 | 488.6 (61%) | 128.6 (49%) | 49.4 | 44.08 | static |
| CB5 | M | 72 | L | 10,574 | 11 | 9.7 | 764.2 (95%) | 229.2 (87%) | 56.5 | 7.39 | double |
| CB6 | F | 68 | R | 64,320 | 26.2 | 16.9 | 193.7 (24%) | 42.2 (16%) | 133.9 | 49.99 | double |
| CB7 | F | 63 | L | 34,123 | 10.6 | 28.3 | 797.4 (99.1%) | 252.6 (96%) | 56.1 | 56.05 | static |
| CB8 | M | 63 | L | 3,707 | 9.1 | 16.2 | 598.3 (74%) | 124 (47%) | 77.2 | 17.64 | double |
| CB9 | F | 52 | R | 22,880 | 10.6 | 30.7 | 798.5 (99.2%) | 254.3 (96%) | 13.2 | 11.77 | static |
| CB10 | M | 63 | L | 9,776 | 237.0 | 46.1 | 703.6 (87%) | 181.8 (69%) | 185.6 | 86.45 | double |
| CB11 | M | 77 | L | 12,500 | 16.6 | 23.2 | 803.5 (99.8%) | 257.9 (98%) | 116.6 | 25.41 | motion |
| N=11 | 5F/6M | 61±13 | 7L/4R | 16.7K±18.6K | 35±68 | 20.5±13 | 601±213 (75%) | 165.4±59 (63%) | 109±85 | 43.2±27 | - |

### Training-induced recovery of visual discrimination and luminance detection sensitivity in CB fields

All patients trained on visual discrimination tasks iteratively, over several months, at multiple blind-field locations (**Fig.1a** and **Table 1**; mean±SD training duration: 20.5±13 months, range 3-46 months), as previously reported (Cavanaugh et al., 2017; Cavanaugh and Huxlin, 2017; Cavanaugh et al., 2015; Das et al., 2014; Huxlin et al., 2009). Chronic CB patients exhibited two types of partially-overlapping improvements: improved discrimination thresholds at trained, blind-field locations (Cavanaugh et al., 2017; Cavanaugh et al., 2015; Das et al., 2014; Huxlin et al., 2009), and recovered luminance detection sensitivity as measured by automated HVF perimetry (Cavanaugh and Huxlin, 2017)–the gold standard, clinical method of assessing visual-field defects. Before training, CB patients were unable to discriminate the orientation (accuracy: 54.6±7.1%) or motion direction (normalized direction range (NDR) thresholds: 96.9±5.8%) of visual targets presented within their blind field (**Fig.1**). This was despite intact performance at corresponding locations within their intact field (intact_*pre*_ *vs* blind_*pre*_; orientation discrimination: t_(9)_=11.5, *p<*.*001*, d=3.63, **Fig.1b**; NDR thresholds: t_(6)_=21.8, *p<*.*001*, d=8.24, **Fig.1c**). Following training, performance improved at trained locations (blind_*pre*_ *vs* blind_*post*_; orientation discrimination: t_(9)_=12.34, *p<*.*001*, d=3.90, **Fig.1b**; NDR thresholds: t_(6)_=18.15, *p<*.*0001*, d=6.86, **Fig.1c**), reaching levels not significantly different from those at intact-field locations (intact_*post*_ *vs* blind_*post*_; orientation discrimination: t_(9)_=1.69, *p=*.*126*, d=0.53, **Fig.1b**; NDR thresholds: t_(6)_=1.80, *p=*.*122*, d=0.68, **Fig.1c**).

**Figure 1.**
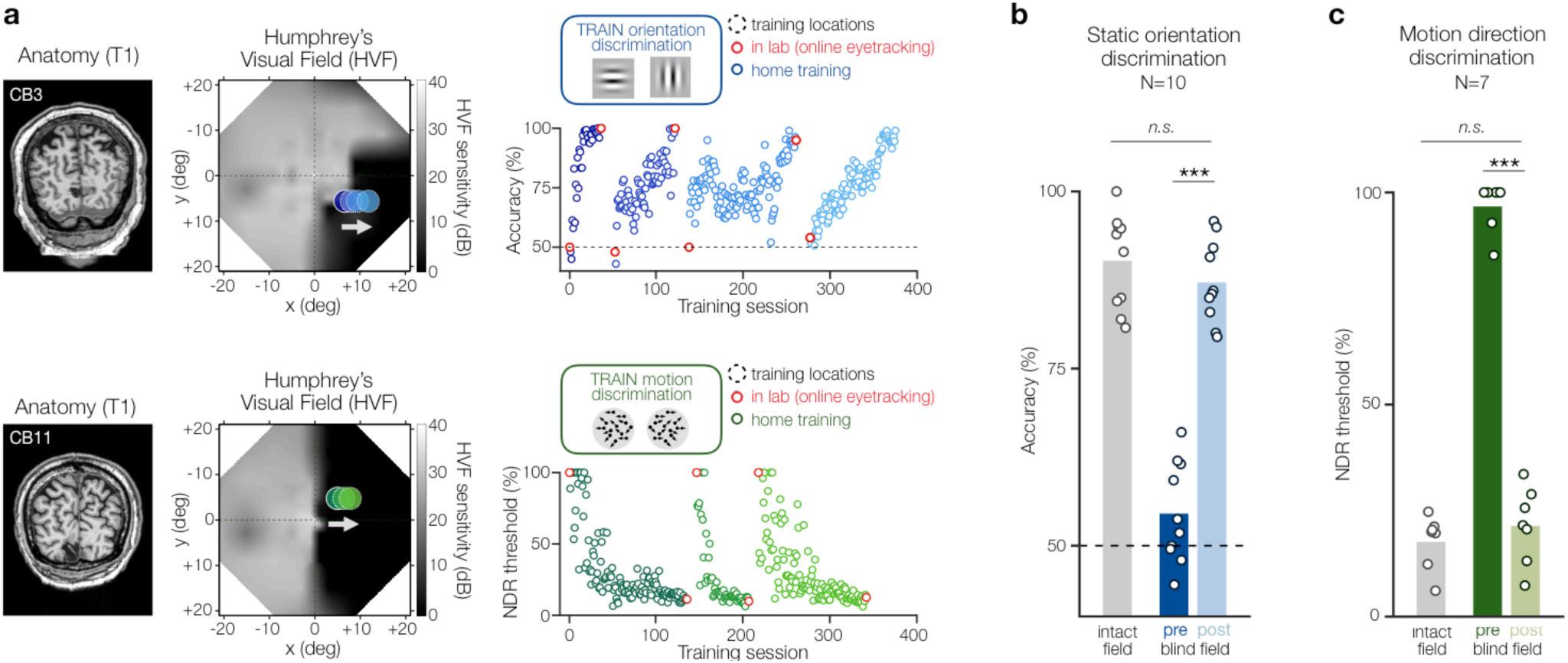
Visual discrimination training recovers visual functions in chronic cortically-blind (CB) fields. **(a)** T1-weighted MRI and 20deg Humphrey Visual Field (HVF) for two of the chronic CB patients illustrating stroke-induced damage to the primary visual cortex and associated loss of conscious luminance detection sensitivity. Dark regions (low HVF sensitivity) correspond to blind-field areas. **(b)** Visual training can successfully restore static orientation discrimination and coarse (left/right) global motion direction discrimination in the blind field of chronic CB patients. Recovery typically requires weeks of daily home training (blue/green dots in scatter plots), is retinotopically-specific, and can be verified in lab under eye-tracking control (red dots in scatter plots). **(c,d)** As detailed in previous work (Cavanaugh et al., 2017; Cavanaugh and Huxlin, 2017; Cavanaugh et al., 2015; Das et al., 2014; Huxlin et al., 2009), all 11 CB patients included in the present study trained on (c) static orientation discrimination (N=10) and/or (d) global motion direction discrimination (N=7) at different blind-field locations. All CB patients recovered performance levels on these tasks similar to those at equivalent locations in their intact hemifields of vision. NDR: normalized direction range of moving dots used in the global direction discrimination task. Bars show mean performance across subjects, with individual data points superimposed.

Critically for the present study, and as recently reported for a larger cohort of CB patients (Cavanaugh and Huxlin, 2017), visual discrimination training did not solely recover performance on trained tasks at trained blind-field locations; it also improved conscious luminance detection HVF sensitivity across both trained and untrained blind-field regions (Cavanaugh and Huxlin, 2017) (**Fig.2** and **Fig.S1**). CB fields are characterized by an abrupt fall-off in HVF sensitivity, from typically-intact (25-30dB) to perimetrically-blind (0-6 dB) levels of HVF sensitivity, with 6dB corresponding to double the test-retest variability of HVF measurements (STATPAC, Zeiss Humphrey Systems). For each patient, the blind-field border delimited impaired regions where the binocular average HVF sensitivity dropped below at least 15dB in the initial (pre-training) HVF map (see Methods). Among the patients included here, the pre-training size of homonymous visual-field defects was 601±213deg^2^ (∼75% of an entire hemifield), ranging from 194 (24%) to 804deg^2^ (99.8%). In the central ±11.5deg (i.e., the restricted visual-field extent stimulated during fMRI), HVF deficits covered an area 165.4±74deg^2^ (63% of the central ±11.5deg hemifield), ranging from 42 (16%) to 258deg^2^ (98%). Following training, all CB patients showed recovery in HVF sensitivity within their blind field (Cavanaugh and Huxlin, 2017) (**Table 1**). As in our previous study (Cavanaugh and Huxlin, 2017), a change ≥+6dB was used as the criterion for HVF recovery (i.e., double the test-retest variability of HVF measurements). We found that HVF sensitivity improved by ≥+6dB over 109±85deg^2^ overall (range: 49-318deg^2^; **Fig.S1**). Within the central 11.5deg (**Fig.2**), improvement occurred over 43.2±27deg^2^ (range: 7.4-86.5deg^2^). Little or no worsening (sensitivity decrements ≥-6 dB) was observed in trained CBs (full HVFs: -2±6.2deg^2^; inner ±11.5deg: -0.9±2.6 deg^2^). The amount of HVF recovery in trained chronic CB is directly proportional to the numbers of training sessions and trained locations (Cavanaugh and Huxlin, 2017), whereas untrained chronic CB patients generally show worsening HVFs over time (Cavanaugh and Huxlin, 2017; Zhang et al., 2006a, b).

**Figure 2.**
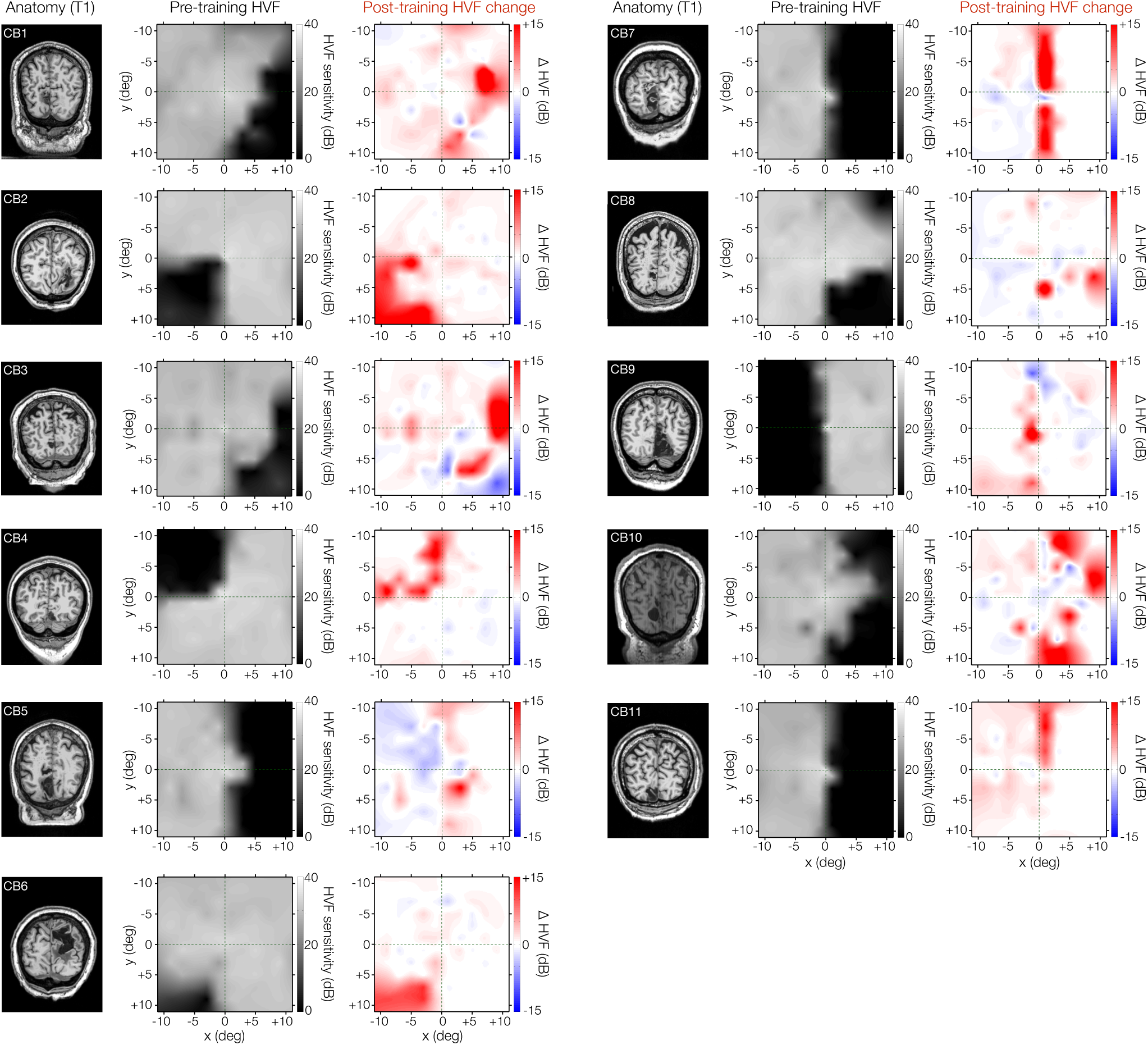
Visual discrimination training improves conscious luminance detection sensitivity in chronic CB patients. T1-weighted MRI and corresponding baseline (pre-training) composite Humphrey Visual Fields (HVF; luminance detection sensitivity in dB) for all 11 chronic CB patients over the central 11.5 degrees of the visual field–i.e., the area stimulated during fMRI retinotopic mapping (see **Fig.S1** for individual maps over full, 24-2 and 10-2 HVFs). Prior to training, all CB patients showed homonymous loss of HVF sensitivity (dark regions) within parts of their visual field. Following training, as indicated by HVF Change maps, all patients showed improved HVF sensitivity (red), which was greatest within the confines of their original blind-field border. A substantial amount of this recovery occurred within the inner 11.5 degrees of the visual field (see (Cavanaugh and Huxlin, 2017) for full description of training-induced HVF recovery in a larger cohort of chronic CB patients). Areas of sensitivity loss (blue) were also noted in some patients, but only 2 (CB3 and CB9) exhibited significant loss (≥-6dB) in their blind hemifields, along with larger HVF improvements.

To better understand the underlying mechanisms of training-induced recovery in luminance detection exhibited by chronic CBs, we compared changes in HVF sensitivity to retinotopic fMRI responses in damaged V1 before and following training.

### Pre-training, spared V1 representations of the blind-field mediate recovery in HVF sensitivity

Qualitatively, intact hemispheres of CB patients and visually-intact controls appeared similar, with V1 and most extrastriate cortical areas (e.g., V2, V3, V4, V3A/B, hMT+) identifiable on retinotopic maps (**Fig.S2**) as in previous studies (Engel et al., 1994; Larsson and Heeger, 2006). The stroke-affected hemispheres of CB patients retained coarse retinotopic organization (**Fig.3a** and **Fig.S3**). V1 was identified in all damaged hemispheres, using both phase reversals and anatomical markers (e.g., spared segments of the calcarine sulcus, comparison to the intact hemisphere). Consistent with prior studies (Kleiser et al., 2001; Morland et al., 2004; Papanikolaou et al., 2014), the visual system of chronic CB patients retained visual representations of the blind field. Before training, a substantial portion of V1 voxels (33±28%) in perilesional cortex of the damaged hemispheres represented parts of the impaired (≤15dB HVF sensitivity) visual field (**Fig.3b,c**). Most of these voxels (24.4±24.1%) corresponded to perimetrically-blind regions (≤6dB HVF sensitivity)–i.e., locations with less than double the test-retest variability of HVF measurements. Such preserved blind-field representations within the damaged V1 have been speculated to be a potential neural substrate for training-induced recovery (Cavanaugh and Huxlin, 2017; Das and Huxlin, 2010; Papanikolaou et al., 2014; Smirnakis, 2016). Here, we directly tested this hypothesis. To do so, we first restricted our analyses to V1 voxels from the damaged hemisphere that represented perimetrically-blind (0-6dB pre-training HVF sensitivity) visual-field locations (**Fig.4a**).

**Figure 3.**
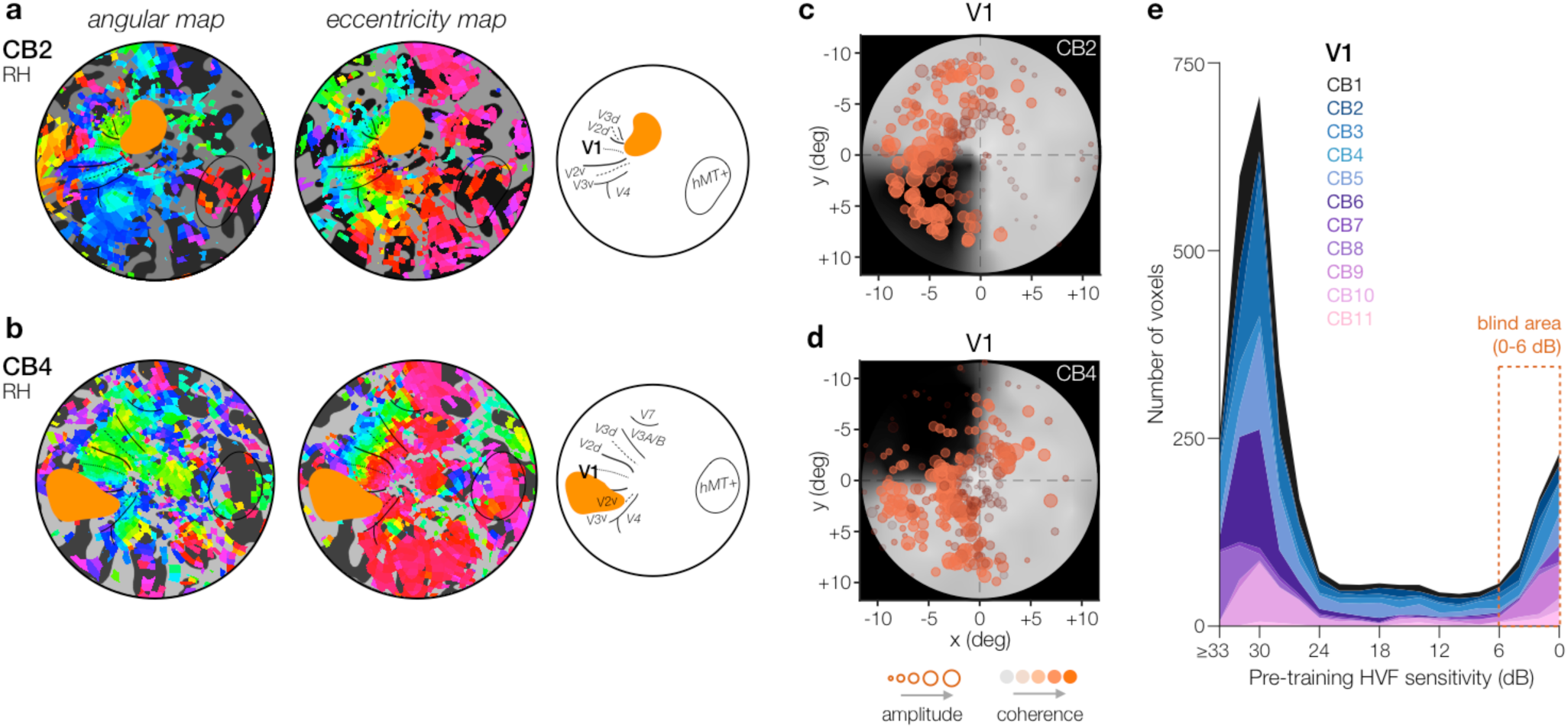
Spared V1 activity representing blind-field areas in chronic CB patients prior to training. **(a,b)** Sample pre-training retinotopic maps of the damaged hemispheres for 2 of our CB patients (see **Fig.S3** for all individual maps). All patients retained retinotopic organization, both in terms of radial and eccentric representations. Stroke-induced lesions (orange mask) overlapped with V1 and other extrastriate areas in most cases, with suggestions of grey matter damages in some patients (e.g., ventricular enlargement, HVF deficits despite no overlap with V1 and extrastriate areas). **(c,d)** V1 voxels from the damaged hemispheres plotted on top of the pre-training HVF (11.5deg) maps for the 2 patients in panel (a,b). Each voxel correspond to an orange circle, with the size denoting response amplitude and depth of orange shading denoting coherence. **(e)** Distribution plot of the number of V1 voxels in the damaged hemispheres of individual chronic CB patients (different subjects=shades from pink to blue) showing a substantial proportion of V1 voxels representing perimetrically-blind regions of the visual field (using 2±1dB steps).

**Figure 4.**
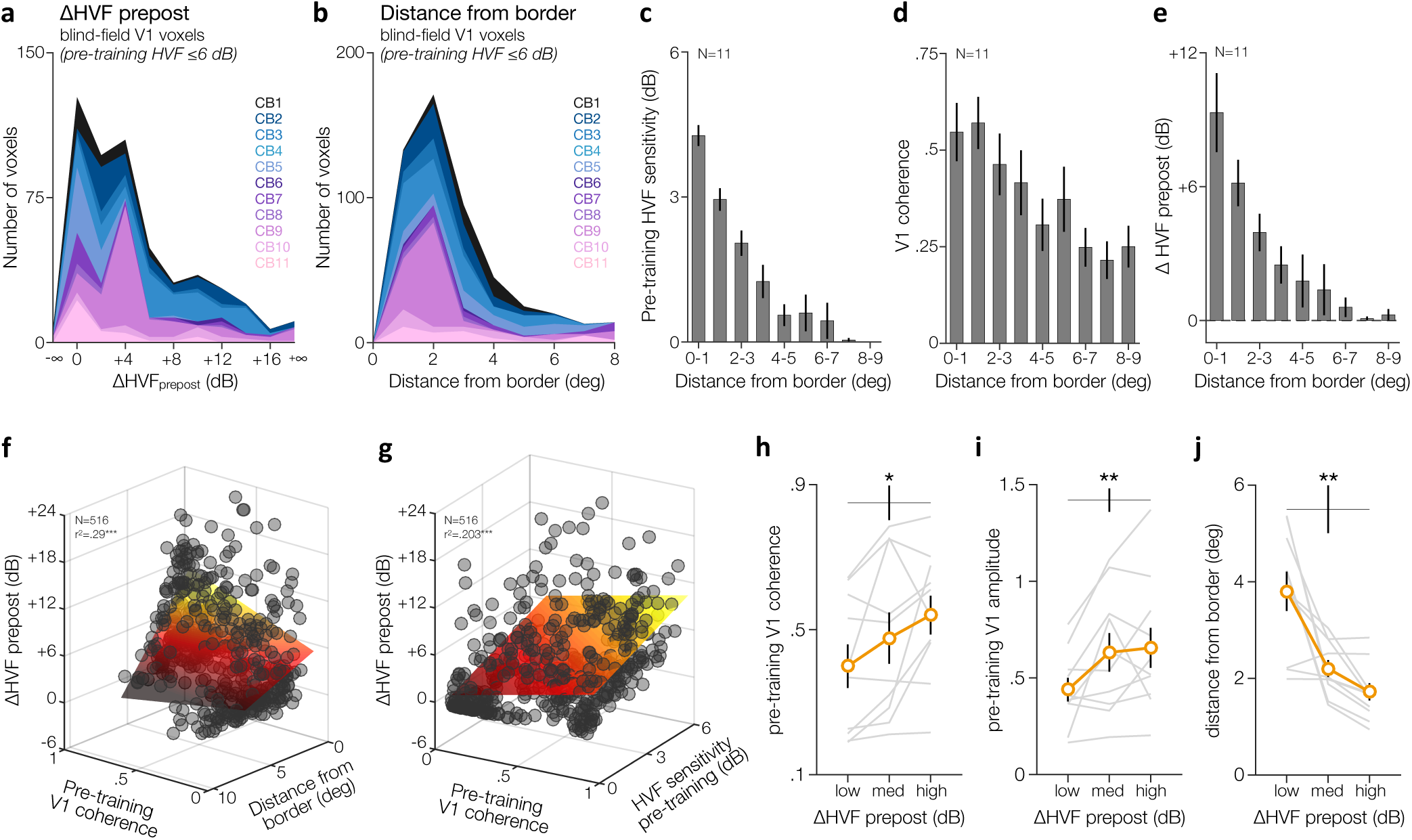
Spared pre-training V1 activity predicts post-training HVF recovery in the blind field of chronic CB patients. **(a,b)** Distribution plot of V1 voxels in damaged hemispheres representing perimetrically-blind regions (≤6 dB HVF sensitivity) prior to training as a function of either (a) the amount of post-training HVF change (using 2±1dB steps), or (b) the distance from the blind-field border (using 1±0.5deg steps). Color conventions as in (Fig.3e). Note the preferential location of these voxels near the blind-field border. **(c-e)** Consistent with enhanced plastic potential along the blind-field border of chronic CB patients, spared blind-field locations closer to the blind-field border showed (c) residual HVF sensitivity, (d) stronger visually-evoked responses, and critically (e) greater HVF change (in dB of sensitivity) from pre-to post-training. Bars represent average estimates across CB patients (±1SEM). **(f,g)** The coherence strength of pre-training V1 voxels was predictive of the magnitude of post-training HVF recovery, with greater HVF recovery occurring (c) closer to the blind-field border, where (d) pre-training HVF sensitivity is not yet completely null. Each gray dot corresponds to a V1 voxel, which were fit using a multiple linear regression model (colored surface). **(h,j)** Pre-training response (h: coherence; i: amplitude) of V1 voxels representing blind-field locations computed for each observer as a function of the level of HVF recovery observed following training (low recovery: ΔHVF_prepost_ ≤+1.83dB; medium recovery: +1.83dB<ΔHVF_prepost_<+5.55dB; high (significant) recovery: ΔHVF_prepost_≥+5.55dB). Error bars correspond to ±1SEM, with gray lines representing individual subjects. **(j)** Same as (h,i) but for distance from the CB border.

As shown in **Fig.4b**, the vast majority of V1 voxels representing portions of the blind field were in close proximity to the initial (pre-training) blind-field border. As expected, pre-training residual HVF sensitivity at these blind-field locations rapidly declined as we moved deeper into the blind field (**Fig.4c**). A similar decrease in V1 visually-evoked response was observed for spared V1 voxels representing blind-field locations further away from the blind-field border (**Fig.4d**). Importantly, most training-induced HVF improvements occurred at blind-field locations represented by V1 voxels near the blind-field border (**Fig.4e**), which further supports the notion (Cavanaugh and Huxlin, 2017) that this may indeed be a region of enhanced plastic potential, where repetitive stimulation provided by training, several months or even years post-stroke, can more readily recover luminance detection sensitivity.

Next, we used a multiple linear regression model across all spared, blind-field V1 voxels (N=516) to assess the relationship between the magnitude of post-training HVF recovery and the strength of pre-training blind-field V1 responses, as a function of the distance from the blind-field border. We found that V1 voxels with stronger visually-evoked responses and representing blind-field regions in close proximity to the blind-field border showed the greatest improvements in HVF recovery following training (**Fig.4f**; F(513,516)=65.4, *p<0*.*0001*; r^2^=.203). A similar result was observed using the level of pre-training residual HVF sensitivity (HVF_pre_; 0-6 dB) rather than the distance from the blind-field border, except that there was no significant interaction term with the strength in pre-training V1 coherence (**Fig.4g**; F(513,516)=65.4, *p<0*.*0001*; r^2^=.203). Finally, the same pattern was also found when using pre-training response amplitude as a measure of the level of pre-training V1 activity at blind-field locations.

The relationship between pre-training V1 activity, distance from the blind-field border, and post-training HVF recovery was present in most CB patients (**Fig.4h-j**). Individually, V1 voxels representing the blind field prior to training were divided into tertiles based on the amount of post-training HVF change (ΔHVF_prepost_) observed across the V1 voxels from all observers. The three separate levels of HVF recovery corresponded to: no recovery (ΔHVF_prepost_≤+1.83dB), moderate recovery: (+1.83dB< ΔHVF_prepost_ ≤+5.55dB), and significant recovery (ΔHVF_prepost_>+5.55dB). This corresponded to V1 voxels representing locations with an average ΔHVF_prepost_ of +0.27±0.64dB (no recovery), +3.27±0.59dB (moderate recovery), or +10.14±1.7dB (significant recovery). Note that CB6 was excluded from this analysis as all V1 voxels within her blind field corresponded to regions of significant HVF recovery, which was mainly due to our restricted access to blind-field areas within the central 11.5 deg (**Fig.2**). For each observer, we computed the mean pre-training V1 activity (i.e., coherence and amplitude of phase-encoded responses). We found that pre-training V1 activity was significantly higher at blind-field locations that exhibited greater HVF recovery following training, both in terms of coherence (**Fig.4h**; F(2,18)=5.16, *p=0*.*017*, η_p_ ^2^=0.36; *no recovery vs. significant recovery*, t(9)=3.10, *p*=0.013, d=.98) and amplitude (**Fig.4i**; F(2,18)=6.32, *p=0*.*008*, η_p_ ^2^=0.41; *no recovery vs. significant recovery*, t(9)=3.67, *p=0*.*005*, d=1.16). Consistent with the fact that training-induced recovery dropped rapidly deeper into the blind field, greater post-training recovery was associated with V1 voxels representing blind-field locations closer to the blind field border (**Fig.4j;** F(1.12,10.05)=15.41, *p=0*.*002*, η_p_ ^2^=0.63; *no recovery vs. significant recovery*, t(9)=4.27, *p=0*.*002*, d=1.16) and with some residual HVF sensitivity (F(2,18)=4.75, *p=0*.*022*, η_p_ ^2^=0.35; *no recovery vs. significant recovery*, t(9)=2.42, *p=0*.*039*, d=.77). Similar patterns of results were observed when dividing V1 voxels based on the strength of pre-training visually-evoked responses, as well as when a more liberal criterion was used to select V1 voxels representing impaired HVF locations (i.e., using 0-15dB instead of 0-6dB).

Taken together, these results provide evidence of enhanced plastic potential near the blind field border of CB patients. They show that the strength of spared V1 activity representing perimetrically-blind areas close to that border predicts recovery of conscious luminance detection sensitivity following training.

### Post-training changes in spared V1 representations of the blind-field

Of the 11 CB patients scanned pre-training, only 8 were able to complete the post-training fMRI session (see *Methods*). Overall, pre- and post-training retinotopic maps were qualitatively similar, suggesting no global reorganization following training (**Fig.5a**; see **Fig.S3,S4** for individual maps). To evaluate changes in retinotopic V1 representations associated with training-induced HVF recovery, we compared mean visually-evoked responses of V1 voxels before and after training, as a function of whether these blind-field locations showed significant recovery or not. For each observer, we compared pre- and post-training mean V1 responses for voxels representing blind-field locations, as a function of the degree of HVF recovery observed at those locations. Voxels were divided into tertiles (defined across both sessions): no/low recovery (ΔHVF_prepost_≤+1.25dB), moderate recovery: (+1.25dB< ΔHVF_prepost_ ≤+5.34dB), or significant recovery (ΔHVF_prepost_>+5.34dB). Training did not affect V1 response coherence, nor interacted with the level of HVF recovery (**Fig.5b**; *all p-values >*.*1*). With respect to response amplitude (**Fig.5c**), there was a main increase in V1 response amplitude with training (F(1,6)=8.43, *p=0*.*027*, η_p_ ^2^=0.58). This significant increase in response amplitude seemed more pronounced for V1 voxels representing blind-field locations with weak pre-training visually-evoked V1 activity and low/no post-training HVF recovery. However, we did not observe a significant interaction between the effect of training and the degree of HVF recovery (F(1,6)=2.53, *p=0*.*121*, η_p_ ^2^=0.30). No global difference in response coherence or amplitude was observed for V1 voxels of the intact hemisphere. In summary, training-induced HVF recovery occurred at blind-field locations with strong, pre-training V1 activity, which did not change further following training. If anything, training enhanced visually-evoked V1 responses for blind-field locations with the weakest, pre-training activity, without an associated, significant increase in visual sensitivity.

**Figure 5.**
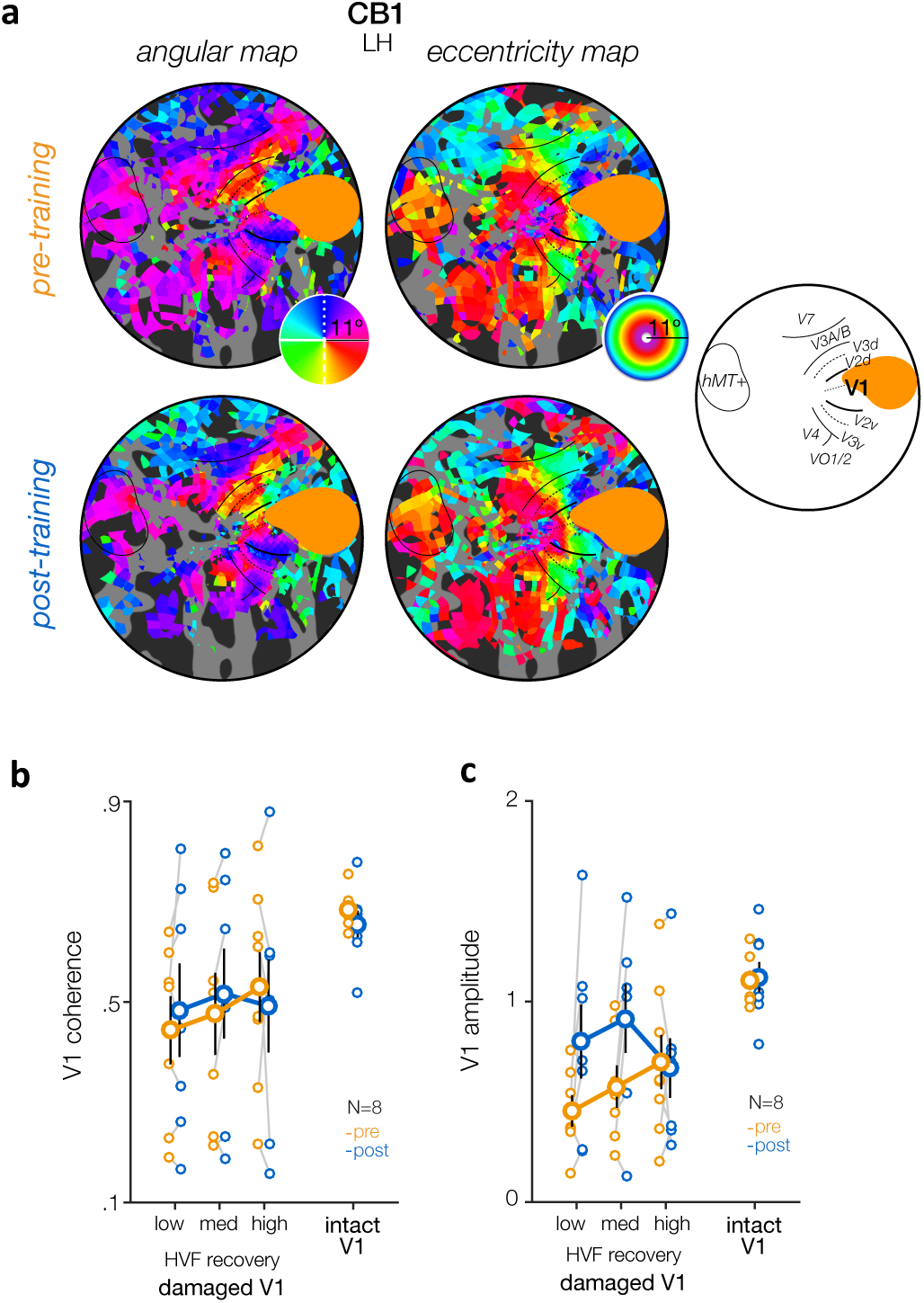
Post-training retinotopic organization and V1 activity following training. **(a)** Sample pre-training and post-training retinotopic maps of the damaged hemispheres for one of our CB patients (see Fig.S3,S4 for individual maps). Qualitatively, no global change in retinotopic organization was observed following training. **(b,c)** Mean V1 response (b: coherence; c: amplitude) for voxels representing blind-field locations prior to training, as a function of training and the amount of post-training HVF recovery (low: ΔHVF_prepost_ ≤+1.25dB; medium: +1.25dB< ΔHVF_prepost_ <+5.34dB; high: ΔHVF_prepost_ ≥+5.34dB). Solid symbols represent group-averaged values with ±1SEM error bars, while each individual CB patient represented by thinner lines representing the pre-post changes at each degree of HVF recovery. No consistent change in V1 coherence was observed following training, regardless of the amount of HVF recovery observed at these blind-field locations. An increase in response amplitude was found, which seemed to preferentially occur at blind-field locations with weak pre-training V1 response that showed limited HVF recovery.

### Increased V1 coverage of the blind field following training

To further characterize changes in V1 associated with training-induced HVF recovery, we used the population receptive field (pRF) method (Dumoulin and Wandell, 2008) to estimate the position and size of the visual-field area that best explained each voxel’s visually-evoked response. For each observer, we assessed how pRF properties changed with training, as a function of whether V1 pRFs covered regions with impaired pre-training HVF sensitivity, or covered only regions with preserved HVF sensitivity (**Fig.6a**; see Methods), excluding pRFs with less than 10% variance explained (r^2^). Consistent with the presence of spared phase-encoded V1 responses within the blind-field (**Fig.3,4**), a substantial number of V1 pRFs (16.3%) were covering blind-field regions prior to training, providing further evidence that perilesional V1 cortex in chronic CB patients retains spared representations of the blind field (Kleiser et al., 2001; Morland et al., 2004; Papanikolaou et al., 2014). Two-way repeated-measures ANOVAs on pRF estimates (**Fig.6b-e**) indicated no effect of training (pre-*vs*. post-training), or interaction between training and HVF sensitivity (impaired HVF *vs*. preserved HVF) on preferred eccentricity, r^2^, or the number of V1 pRFs (all *p>0*.*1*). As expected, there were fewer V1 pRFs covering blind-field regions than pRFs covering visual-field locations with preserved HVF sensitivity (F(1,7)=29.9, *p<0*.*001*, η_p_ ^2^=0.81). Blind-field pRFs had a marginally lower variance explained (F(1,7)=4.95, *p=0*.*061*, η_p_ ^2^=0.41), and were both more eccentric (F(1,7)=28.88, *p<0*.*001*, η_p_ ^2^=0.81) and larger (F(1,7)=33.85, *p<0*.*001*, η_p_ ^2^=0.83) on average.

**Figure 6.**
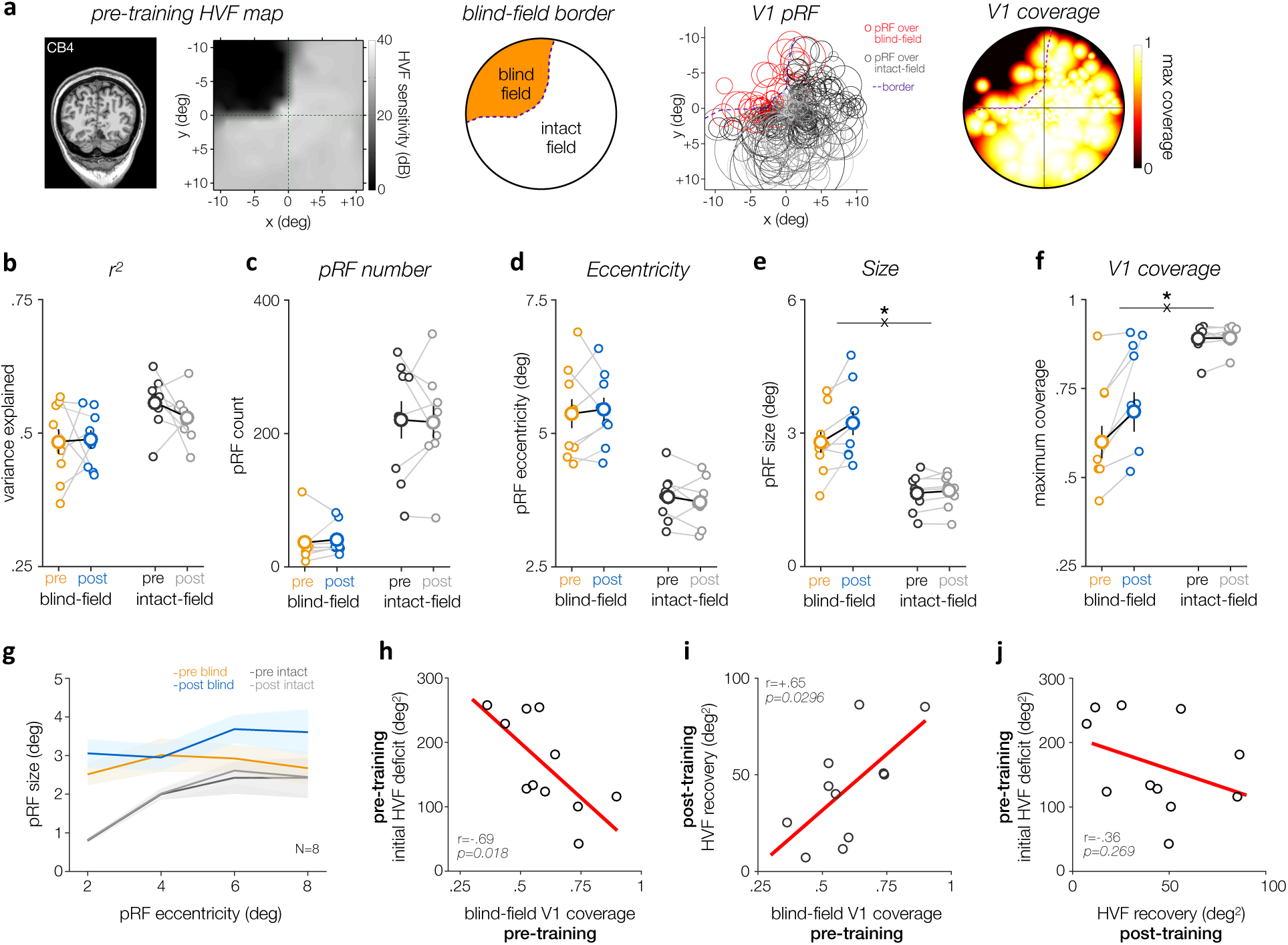
Enhanced population receptive field size and coverage of the blind field in V1 following training. **(a)** The population receptive field method was used to estimate the position and size of the visual-field area that best explained each voxel’s visually-evoked response. The initial (pre-training) HVF map was used to define the blind-field border and determine whether each single pRF covered blind-field regions or solely intact-field regions with preserved HVF sensitivity. **(b-e)** Effect of training on pRF estimates for V1 pRFs covering either intact or blind-field regions. Training only resulted in a significant increase in pRF size for pRF covering blind-field regions. Solid symbols show group-averaged estimates (±1SEM), and smaller dots individual data. **(f)** Chronic CB patients showed enhanced V1 coverage over blind-field regions following training. Same convention then (b-e). **(g)** pRF size plotted as a function of eccentricity indicated larger pRF size in perilesional V1 cortex prior to training. Following training, pRF covering the blind field increased in size across eccentricity. Shaded areas correspond to ±1SEM. **(h)** Pre-training V1 coverage of the blind field correlated with the initial HVF deficit area. **(i)** Pre-training V1 coverage of the blind field was predictive of the amount of HVF recovery observed following training. **(j)** Pre-training HVF deficit area did not correlate with the amount of HVF recovery observed with training.

Critically, we found a significant effect of training on pRF size (F(1,7)=9.41, *p=0*.*018*, η_p_ ^2^=0.57), with a significant interaction between training and HVF sensitivity (F(1,7)=5.88, *p=0*.*045*, η_p_^2^ =0.46). These effects reflect a significant enlargement of V1 pRFs covering the blind field by ∼+17% following training (t(7)=2.86, *p=0*.*024, d=1*.*01*), whereas there was no significant change in size for pRFs covering solely regions with preserved, baseline HVF sensitivity (t(7)=1.00, *p=0*.*349, d=*.*36*). To further characterize this post-training enlargement of V1 pRFs covering the blind field, we estimated pRF size as function of eccentricity (see Methods) (**Fig.6g**). A previous study (Papanikolaou et al., 2014) reported the presence of larger pRF size in perilesional V1 cortex of untrained, chronic CB patients, reflecting the presence of spared V1 representations along the blind-field border. Consistent with this finding, we found that V1 pRFs covering blind-field regions were larger than pRFs covering solely intact-field regions at corresponding eccentricities (F(1,7)=27.74, *p=0*.*001*, η_p_ ^2^=0.80), which interacted with the significant increase in pRF size with eccentricity (eccentricity*HVF sensitivity interaction: F(3,21)=4.43, *p=0*.*015*, η_p_ ^2^=0.39). The increase in pRF size with eccentricity for V1 pRFs covering intact-field regions was similar to visually-intact controls (all F<1). Furthermore, training increased pRF size (F(1,7)=20.54, *p=0*.*003*, η_p_ ^2^=0.75), particularly for V1 pRFs covering the blind field (training*HVF sensitivity interaction: F(1,7)=5.61, *p=0*.*050*, η_p_ ^2^=0.45). This increase in pRF size in proximity of the blind-field border after training was constant across eccentricities (training*eccentricity interaction: F(3,21)=1.67, *p=0*.*204*, η_p_ ^2^=0.19).

Finally, we derived visual-field coverage maps over the central 11.5 deg by superimposing all V1 pRFs and computing the coverage index for regions with either preserved or impaired HVF sensitivity (**Fig.6a** and *Methods*). Consistent with the post-training enlargement of pRF size over blind-field areas (**Fig.6e**), training increased V1 coverage of the blind-field (**Fig.6f;** training*HVF sensitivity interaction: F(1,7)=10.26, *p=0*.*015*, η_p_ ^2^=0.59). Visual-field locations with preserved HVF sensitivity were well covered by V1 pRFs, with no difference between testing sessions (pre: 88.93±4.2%; post: 89.39±3.2%; t_7_=0.73, *p=0.489* d=.26), and were similar to visually-intact age-matched controls (88.28±2.8%; F(1,15)=0.14, *p=.711*). Prior to training, CB patients showed overlap between pRF V1 maps and perimetrically-blind regions (**Fig.6f**; pre-training coverage index: 62.61±15.22%, range: 43.5-89.7%). Following training, V1 coverage over blind-field areas improved (Wilcoxon signed-rank test: *p=.008*), while no change was found for pRFs representing intact regions of the visual field (*p=.547*). Pre-training V1 coverage was negatively correlated with pre-training HVF deficit area (**Fig.6h**; r=-.693, *p=0.018*). Notably, we found pRF mapping also appeared able to serve as a simple predictor of the recovery potential of chronic CB patients. Indeed, the amount of spared, pre-training V1 coverage over the blind field was significantly correlated with the area of HVF improvement observed after training within the central ±11.5deg of the visual field (**Fig.6i**; r=+.652, *p=0.029*). A similar trend was still observed when computing coverage maps using pRFs weighted by their own variance explained (r=.534, *p=0.0905*). Note that while the pre-training deficit area estimated from HVF maps correlated with the amount of pre-training V1 coverage of the blind field, the area of pre-training HVF deficit was not a good predictor of the amount of HVF improvements observed post-training (**Fig.6j**; r=-.365, *p=0.269*). These results provide further evidence that spared V1 representations within perimetrically-blind regions of the visual field mediate recovery of conscious, luminance-detection sensitivity following training. Moreover, they detail what level of functionality of V1 circuits is needed to support such training-induced improvements. As such, while the increase in blind-field coverage following training did not correlate with the amount of training-induced HVF recovery, we speculate that it likely represents just the increase in V1 responsiveness needed to support further recovery, deeper within the blind field of chronic CB patients, once additional training is administered.

## Discussion

In the present study, we used fMRI retinotopic mapping to study the neural substrates of training-induced recovery of luminance detection sensitivity (HVF) in eleven CB patients with large, chronic, hemianopic visual-field defects due to stroke-induced V1 damage. Six main findings emerged: (1) prior to training, chronic CB patients exhibited substantial, visually-evoked fMRI responses in perilesional V1 corresponding to blind regions of their ipsilesional hemifields; (2) spared, pre-training V1 activity representing blind-field regions in close proximity to the blind-field border was predictive of the amount of luminance detection sensitivity recovered post-training; (3) no further change in V1 activity (coherence or response amplitude) was observed post-training at blind-field locations exhibiting greatest HVF recovery; (4) pre-training V1 coverage of the blind field was also predictive of the amount of training-induced HVF recovery; (5) training enhanced V1 response amplitude at blind-field locations, particularly for locations with weak pre-training V1 activity and limited HVF improvements; (6) training-induced recovery was also associated with increased V1 coverage over blind-field regions, mostly mediated by an enlargement in pRF size. Taken together, our findings provide vital and novel insights regarding the neural mechanisms by which daily visual discrimination training administered over several months restores conscious luminance-detection sensitivity in the blind field of chronic CB patients. Above all, our results provide empirical evidence that spared V1 circuits representing blind-field regions serve as critical substrates of training-induced recovery in chronic CB.

Most prior neuroimaging studies on the properties of cortical reorganization and residual visual processing in CB involved case reports (Baseler et al., 1999; Bridge et al., 2010; Bridge et al., 2008; Dilks et al., 2007; Henriksson et al., 2007; Vaina et al., 2014). Only more recent investigations have examined larger patient cohorts (Ajina et al., 2015; Martin et al., 2012; Papanikolaou et al., 2014; Raemaekers et al., 2011). Moreover, previous studies often faced interpretational issues that arose from significant inter-individual variability in the type, size, and age of cortical damage, among others. Here, we restricted our recruitment to patients with visual impairments resulting from chronic, stroke-induced V1 damage sustained during adulthood. The intent was to minimize inter-subject variability from diversity in underlying pathology (e.g., stroke *versus* resection or trauma), and from changes due to increased plasticity from damage early in development. Like prior work (Baseler et al., 1999; Papanikolaou et al., 2014; Schmid et al., 2010), we found that V1 damage did not induce large-scale reorganization of phase maps. Despite their stroke, all CB patients in the current study showed differentiable V1/V2 borders and phase reversals between most extrastriate visual areas. Importantly, we found preserved, perilesional V1 cortex representing regions of the blind field, consistent with prior studies on smaller groups of CB patients (Kleiser et al., 2001; Morland et al., 2004; Papanikolaou et al., 2014). Such residual visual representations within regions of the blind field in CB patients differ from controls tested with artificial scotomas, who only show limited visually-evoked activity and pRF coverage of masked visual-field locations (Binda et al., 2013; Hummer et al., 2017; Papanikolaou et al., 2014).

The existence of residual, visually-evoked activity representing blind-field regions in the damaged V1 of CB patients has important implications, both for understanding preserved vision and designing better visual restitution strategies (Ajina and Bridge, 2016; Ajina and Kennard, 2012; Chokron et al., 2008; Das and Huxlin, 2010; Herpich et al., 2019; Morland et al., 2004; Papanikolaou et al., 2014; Perez and Chokron, 2014; Smirnakis, 2016; Wandell and Smirnakis, 2009). Although it is unclear why regions with preserved V1 activity do not mediate conscious perception without training, here we reveal the potential relevance of such preserved activity to visual restoration efforts. Specifically, the strength in pre-training fMRI activity representing blind-field regions in the damaged V1, as well as the amount of pre-training V1 coverage of the blind field, were predictive of the magnitude of training-induced HVF recovery. This finding indicates that pre-training retinotopic mapping can provide vital insights into a chronic patient’s potential for training-induced recovery; indeed, future, more efficient rehabilitation strategies could optimize training by pre-selecting regions more amenable to recovery.

Another key finding here was that blind-field locations with the greatest HVF recovery did not exhibit further increases in the strength of their visually-evoked BOLD responses post-training. This suggests that training-induced perceptual recovery was mediated by improvements in the read-out efficiency of sensory signals at blind-field locations with strong, initial (pre-training) V1 activity. Such a learning mechanism would be consistent with observations in intact visual systems, where improved behavioral performance with training often involves changes in sensory read-out efficiency at later decoding stages, rather than in early sensory representations (Chowdhury and DeAngelis, 2008; Kumano and Uka, 2013; Law and Gold, 2008). Of special relevance given our patients’ training regime, training-induced improvements in motion discrimination in intact (non-human primate) visual systems are not associated with enhanced firing rates in the motion-sensitive area MT, but with changes in firing rates in the lateral intraparietal area (LIP), which is implicated in accumulating sensory evidence (Law and Gold, 2008). Moreover, we previously showed using a drift diffusion model of decision making applied to fMRI responses, that untrained CB patients suffer from slower rate of information accumulation (i.e., reduced drift rate) when performing a global direction discrimination task at blind-field locations, relative to their intact fields and to intact controls (Martin et al., 2012). Importantly, untrained chronic CB patients do not exhibit spontaneous visual recovery, either with respect to motion perception or luminance detection (i.e. HVF perimetry)(Cavanaugh and Huxlin, 2017; Zhang et al., 2006a, b). Thus, the mere presence of spared V1 activity representing portions of the blind-field is not sufficient for recovery to occur; deliberate, localized training is required to recover conscious luminance detection within chronic CB fields. Although these results for the largest improvements in luminance detection sensitivity suggest plasticity at neural decoding rather than encoding stages, we did find some evidence suggesting post-training changes in V1 representations, particularly for areas with no HVF recovery. These blind-field locations, represented by weak pre-training V1 activity, showed increased response amplitudes and pRF coverage post-training. We speculate that a certain level of baseline V1 responsiveness is necessary for training to effectively induce recovery of conscious luminance detection. In this context, locations that remained perimetrically blind but showed stronger post-training V1 activity might serve as a new *baseline* upon which further training could act to induce additional HVF recovery–consistent with the observation that sequential, iterative training can recover vision progressively deeper within chronic CB fields (Cavanaugh and Huxlin, 2017; Das et al., 2014; Huxlin et al., 2009).

Training-induced HVF recovery was also associated with larger pRF size at the blind-field border, increasing V1 coverage of the blind field post-training. This finding is consistent with single-unit electrophysiological data in lesioned animals showing RF enlargement at the border of V1 lesions following repetitive stimulation of unresponsive locations (Schweigart and Eysel, 2002). Enhanced plasticity in perilesional V1 cortex is also consistent with the fact that training-induced HVF improvements are generally concentrated along the blind-field border of CB patients (Cavanaugh and Huxlin, 2017). It is unclear whether and how visual training affects pRF properties, both in neurotypical and patient populations. Several studies have reported changes in the position and size of RF (Womelsdorf et al., 2006) and pRF (Klein et al., 2014) when spatial attention is covertly deployed to a peripheral location. Our visual rehabilitation approach required patients to covertly attend to specific regions just inside their blind-field border, which could induce increased blind-field coverage during training. With repetitive training, changes in pRF properties may become permanent and account for increased pRF size and blind-field coverage we observed in V1. Although we did not test untrained, chronic CB patients, such patients do not exhibit spontaneous HVF recovery (Cavanaugh and Huxlin, 2017; Zhang et al., 2006a, b). HVF recovery in trained CB patients correlates with the amount of training performed, while untrained chronic CB patients only show restricted HVF improvements counterbalanced by areas of HVF worsening (Cavanaugh and Huxlin, 2017). Thus, limited changes in V1 activity, if any, would be observed in untrained, chronic CB patients.

The fundamental role of spared V1 cortex in visual rehabilitation does not rule out the contribution of extrastriate cortex to this process. Our results show that a certain level of spared V1 activity is necessary but not sufficient for training-induced HVF recovery, which likely involves feedback from extrastriate cortex as typically observed in intact neural systems during training (Li, 2016). Moreover, while HVF recovery might rely on V1, trained CB patients can also recover relatively complex visual functions (Das et al., 2014; Huxlin et al., 2009; Melnick et al., 2016b), which presumably require processing by higher-level areas. For instance, recovery of motion direction discrimination may rely primarily on extrastriate motion-selective areas (e.g., hMT+)(Hagan et al., 2019; Melnick et al., 2016a). Given the dramatic impact of V1 damage on most, if not all, visual functions, and the diversity of stimuli whose processing can be restored in CB (Chokron et al., 2008; Melnick et al., 2016b), the road to full recovery likely depends on recruitment of and interactions between multiple visual areas. The role of extrastriate areas might be better characterized by measuring brain activity while CB patients are asked to discriminate visual information within their blind field, and at different training stages.

To conclude, the present study reveals a hitherto unrecognized and potentially vital role of perilesional V1 cortex in mediating training-induced recovery of conscious luminance detection sensitivity in chronic, stroke-induced CB. Our findings indicate that recovery relies on pre-training, spared, V1 activity representing blind-field locations. Moreover, training increased V1 coverage over blind-field regions, which was mediated by the enlargement of V1 pRFs in close proximity to the initial blind-field border. Aside from improving our scientific understanding of the neural mechanisms underlying visual recovery in CB, our findings are of vital importance for the development of more principled, customized, clinical neurorehabilitation therapies, adapted to the increasing number of people who suffer from cortically-induced visual impairments. Targeting blind-field regions with spared V1 activity might substantially improve the benefits of visual restoration training, particularly in subacute post-stroke periods (i.e., first 3 months) during which our training paradigm can substantially enhance capacity for recovery and preserve visual functions (Saionz et al., 2019).

## Acknowledgments

The present study was funded by NIH (EY027314 and EY021209 to K.R.H.; Core Center Grant P30 EY001319 and training grant T32 EY007125 to the Center for Visual Science; a pre-doctoral NRSA EY025918 to MRC), and an unrestricted grant from the Research to Prevent Blindness (RPB) Foundation to the Flaum Eye Institute. The authors thank Terrance Schaefer, who performed Humphrey visual field tests and to Patricia Weber for running the MRI on all patients presented here. We also thank Drs. Steven Feldon, Zoe Williams, Bogachan Sahin, Ania Busza, Shobha Boghani, Alexander Hartmann and Ronald Plotnick for referring CB patients to us for study.

## Author contribution

AB, AD, EPM DJH and KRH designed the study; AB, MDM, MRC and AD collected the data; AB, MDM and EPM analyzed the data; AB and KRH wrote the manuscript, and all authors commented on it.

## Data availability

All data are available from the authors upon request.

## Methods

### Participants

Eleven adult CB subjects (5 females; see **Table 1** for demographics), ranging in age from 29 to 77 years old (mean: 61±13 years old) underwent visual retraining starting at least 5 months (35±68 months, ranging from 5 to 237 months) after stroke-induced occipital damage (verified from structural MRI; **Fig.2** and **Fig.S1**). These patients had large, contralesional, homonymous visual field defects determined from 24-2 and 10-2 Humphrey Visual Fields (HVFs) and shown in **Fig.2 and Fig.S1**. Recruited subjects had no ocular health problems, neurological and/or cognitive impairments, and none of the subjects suffered from visual or other forms of neglect. All 11 CB participants were scanned pre-training, but only 8 could complete the post-training fMRI session (CB1-8). CB10 had become MRI-incompatible at the time of the post-training visit, and technical difficulties with the magnet meant that post-training MRI could not be collected at the time of CB9 and CB11’s post-training visits. Nine visually- and neurologically-intact subjects (7 females, mean age: 50±14 years, range: 28 to 65 years) served as age-matched controls. The University of Rochester’s Research Subjects Review Board approved all experiments conducted as part of this study. The experimental procedures adhered to the tenets of the Declaration of Helsinki and were conducted after obtaining the subjects’ informed, written consent.

### Visual discrimination testing and training

CB subjects were trained at select blind-field locations on coarse (left-right) direction discrimination of random dot stimuli and/or vertical-horizontal orientation discrimination of static Gabor patches (**Table 1** and **Fig.1**). For details about the training, testing stimuli and tasks, see our published studies (Cavanaugh et al., 2017; Cavanaugh and Huxlin, 2017; Cavanaugh et al., 2015; Das et al., 2014; Huxlin et al., 2009; Saionz et al., 2019). Training was started at visual-field locations where discrimination performance on either task first dropped to chance (50% correct) during blind-field border mapping (i.e., in lab with online fixation monitoring using an infrared eye tracker [Eyelink 1000 or ISCAN RK464] interfaced with our psychophysical software, to ensure gaze-contingent stimulus presentation). Patients then trained at home, performing 300 trials per day, per location, at least five days per week. Progress was assessed via weekly analysis of data log files automatically generated by our training software and sent to the lab. Once performance increased to a level comparable to that at equivalent, intact-field locations (measured during pre-testing), the training location was moved 1° deeper into the blind field along the x-axis. While home training was performed without eye-tracking, patients were repeatedly instructed to fixate the central fixation spot. Critically, all performance on the trained tasks was verified with online eye-tracking in lab during the post-training visit, and patients were not considered to have recovered unless in-lab performance demonstrated improvement matching that recorded during home training.

### Perimetric visual field tests

Humphrey visual field (HVF) tests (24-2 and 10-2) were performed in all patients using a Humphrey Field Analyzer (HFA II 750), administered by the same ophthalmic technician, at the Flaum Eye Institute of the University of Rochester Medical Center. HVF perimetry measures monocular luminance detection thresholds (in dB) by testing light intensities in a regular grid over the central 48° (24-2 HVF) or 18° (10-2 HVF) of the visual field, with fixation controlled using built-in eyetracking. HVF thresholds (in dB) reflect the extent to which light can be dimmed from the maximum intensity (3,174 cd/m^2^) and still be detected relative to background (∼10 cd/m^2^) at each test location. For all patients enrolled in the present study, percentage of fixation losses, false positive and false negative responses were <20%. Given the homonymous nature of their deficits, composite binocular HVFs were generated by averaging luminance sensitivity from monocular HVFs at each location between both eyes. Then, we interpolated between test locations to obtain smooth composite HVF with 0.1deg^2^ resolution (see our previous study (Cavanaugh and Huxlin, 2017) for more details; see **Fig.2** and **Fig.S1** for individual maps). Several metrics calculated by the Humphrey STATPAC software (Zeiss Humphrey Systems) can be used to define visual defects and quantify HVF changes in CB patients (Cavanaugh and Huxlin, 2017). Here, we specifically measured the area of the HVF in which changes in sensitivity (increase or decrease) occurred. To do so, differences in HVF maps were computed by first subtracting pre- and post-training, non-interpolated HVF maps, and then interpolating HVF sensitivity change (ΔHVF_prepost_) between test locations. CB fields are characterized by an abrupt fall-off in HVF sensitivity, from typically-intact HVF levels (25-30dB) to perimetrically-blind (0-6 dB) levels. Perimetrically-blind areas were defined as areas with less than 6dB HVF sensitivity, which is double the test-retest variability of HVF measurements (STATPAC, Zeiss Humphrey Systems). Similarly, and as in our previous study (Cavanaugh and Huxlin, 2017), an improvement by at least +6dB was used as the criterion for HVF recovery, and a decrease by at least -6dB for HVF worsening. The area of HVF impairment (**Table 1**) was computed as the area of impaired HVF sensitivity (i.e., impaired binocular average HVF sensitivity below 15dB), which was used to delimitate the border of the blind field from the pre-training HVF measurement and compute a map of the distance from the blind-field border.

### MRI acquisition

MRI data were acquired on a 3T Magnetom Trio scanner (Siemens, Erlangen, Germany) using a 32-channel head coil. Functional scans were acquired with gradient recalled echo-planar imaging to measure blood oxygen level-dependent (BOLD) changes in image intensity (Ogawa et al., 1990): 21 slices oriented perpendicular to the calcarine sulcus; repetition time 1.5s; echo time 30-ms; flip angle 75º; voxel size 3×3×3 mm; grid size 64×64. At the beginning of each scanning session, an in-plane T1 weighted (MPRAGE, magnetization prepared rapid gradient echo) anatomical volume was acquired in the same slices as the functional scans, but with voxel size of 0.75×0.75×3 mm. In addition, three high-resolution, T1-weighted anatomical volumes (MPRAGE, 1×1×1 mm) were acquired. The high-resolution T1 volumes were averaged together, and the average was used to extract and computationally flatten the cortical surface using FreeSurfer (http://surfer.nmr.mgh.harvard.edu). In-plane, anatomical images were aligned with the average high-resolution anatomical volume by an automated, robust image registration algorithm (Nestares and Heeger, 2000). The alignment parameters were used to project the measured fMRI responses onto the flattened cortical surfaces for visualization.

### Retinotopic mapping

Travelling wave stimuli were used for retinotopic mapping (**Fig.3** and **Fig.S2,S3**), which consisted of clockwise/counterclockwise rotating wedges and expanding/contracting rings filled with high-contrast, black-and-white sliding checkerboard stimuli. Wedges spanned 0.5-11.25º in radius; i.e., a small gray circle (1º diameter) was in the middle of the checkerboard stimuli, within which appeared a fixation cross. All subjects performed a fixation task during scanning (see below for details). Both the rings and the wedges had a 25% duty cycle. Each run in a session consisted of 10.5 cycles (24 s length) of the stimulus rotating or expanding/contracting (168 volumes). The first 8 volumes (0.5 cycle) were discarded prior to data analysis. Each scanning session consisted of six runs of the wedge stimulus (3 clockwise and 3 counterclockwise) and four runs of the ring stimulus (2 expanding and 2 contracting). The human MT/MST complex (hMT+) was defined by measuring BOLD responses to coherent vs. incoherent motion (block alternation protocol, 18 s period). The motion stimulus consisted a 10° radius circular aperture with moving white dots presented on a black background (average density of 3 dots/deg^2^). Epochs of coherent motion consisted of 100% coherent translational motion (7 deg/s), randomly changing in direction every 3s (6 possible motion directions). During epochs of incoherent motion, each dot was assigned a random direction on every frame of the stimulus. Each full cycle was 18s long (12 frames) and the stimulus was run for 11 cycles (176 volumes). The first 16 volumes (1 cycle) were discarded prior to data analysis.

### Behavioral task during MRI scanning

All subjects performed a task at fixation to minimize fixation breaks with large eye movements and to ensure consistent behavioral and attentional state throughout the fMRI data acquisition. A two-interval forced-choice (2-IFC) luminance decrement detection task was performed with respect to the fixation cross. A cyan cross was first displayed at the beginning of each trial and was briefly dimmed during each interval. The two target intervals were separated by a 500ms period, during which the cross’ color changed back again to cyan. After a final 500ms, the cross turned yellow signaling that the subject should report which of the two intervals contained the dimmer cyan cross by pressing one of 2 buttons. After the button press, the fixation cross changed from yellow to either green for a correct response, or to red for an incorrect response as well as if the subject did not respond. The target luminance decrement was adjusted throughout each scanning session using a 2-down/1-up staircase to maintain performance near 71%-correct and maintain attention at fixation during both pre-training (71±9%) and post-training (73±4%) scanning sessions.

### MRI lesion reconstruction

Lesions were defined by hand on the high-resolution, T1-weighted, anatomical images. Lesions were largely filled with CSF and/or had relatively low image intensity, they were easily delineated on T1 anatomical images. Lesion reconstruction was performed using the FreeSurfer software by filling voxels corresponding to the lesioned area with an image intensity corresponding to that of white matter. These voxels were then added to the white matter mask generated by FreeSurfer, creating a surface boundary at the approximate location in the brain where gray matter had existed prior to the lesion. This procedure minimized distortions in the cortical reconstruction, and facilitated visualizing the lesions on flattened gray matter maps (e.g. orange areas in **Fig.3a,b** and **Fig.5a**; see also **Fig.S3,4** for all individual maps).

### Phase-encoded retinotopic analysis

All fMRI data analysis steps were performed with custom software (mrTools, http://www.cns.nyu.edu/heegerlab) written in Matlab (The MathWorks, Natick, MA). The first half-cycle of each fMRI run was not included in subsequent analysis to allow longitudinal magnetization and hemodynamics to reach steady-state. The data from each run were preprocessed using standard procedures for motion compensation (Jenkinson et al., 2002; Nestares and Heeger, 2000) and detrending(Purdon and Weisskoff, 1998; Zarahn et al., 1997). The data were then analyzed by fitting a sinusoid to the time series of each voxel, and by computing the correlation/coherence between the measured time series and the best-fitting sinusoid (Bandettini et al., 1993; Engel et al., 1997). Data acquired with different stimulus directions were combined to estimate the response phase independent of the phase lag caused by the hemodynamic delay of the fMRI response. Time series data for each run were first coarsely corrected for the hemodynamic delay by shifting the time series of each voxel back three time points (4.5 s). The time series for the counterclockwise wedge and contracting rings were time-reversed and averaged with the time series data for clockwise wedges and expanding rings, respectively. This way, any residual phase lag was canceled allowing us to directly convert response phase into polar angle or eccentricity without having to estimate hemodynamic delay. The two motion localizer runs were directly averaged together. The Fourier transforms of the resulting time series were obtained and the amplitudes and phases at the stimulus frequency were examined. Coherence was computed as the ratio between the amplitude at the stimulus frequency and the square root of the sum of squares of the amplitudes at all frequencies. Coherence, amplitude and phase values from each gray matter voxel were visualized on flat maps of the occipital cortex centered on the occipital pole (**Fig.S2-4**). V1 boundaries, as well as boundaries between both ventral and dorsal extrastriate visual-cortical areas were identified in the flattened retinotopic maps as phase reversals in polar angle components (Engel et al., 1997; Engel et al., 1994; Larsson and Heeger, 2006) and used to draw specific regions-of-interests (ROIs) on the flat maps, before being converted to cortical depth. Similar to previous retinotopic mapping studies (Engel et al., 1994; Larsson and Heeger, 2006), we could identify V1, ventral (V2v, V3v, V4, VO1/2) and dorsal (V2d, V3d, V3A/B) extrastriate cortical areas, as well as the human motion-sensitive complex (hMT+) in intact hemispheres of chronic CB patients and visually-intact controls (**Fig.S2**). To improve the accuracy of retinotopic maps in the damaged hemispheres of CB patients (**Fig.S3**), intact anatomical landmarks and comparisons to the intact hemispheres were used. On rare occasions, some extrastriate areas were not identifiable, due either to a lack of activity, suggesting overlap with the lesion (e.g., CB6), or an absence of clear phase reversals between certain areas (e.g., CB11). Coherence, amplitude and phase values of each voxel within a ROI were used for further ROI analysis. In addition, each voxel was associated to various HVF measures based on its retinotopic position (e.g., pre-training HVF sensitivity, pre-post change in HVF sensitivity, distance from the blind-field border). Voxels with less than 10% coherence were excluded from further analyses.

### Population receptive field (pRF) analyses and coverage maps

The pRF method (Dumoulin and Wandell, 2008; Wandell and Winawer, 2015) estimates the position and size of each voxel’s RF by determining the area of the visual field that best explains the relation between the time-course of the retinotopic mapping stimulus and the measured BOLD time series. The pRF model implements an isotropic Gaussian spatial receptive field, whose center and radius are derived by fitting the voxel’s BOLD signal responses to the model’s estimated responses elicited by convolving the model with the retinotopic mapping stimuli. pRF depict the area of the visual field that effectively evokes a response for a given voxel. pRF centers were limited to 0.5-10deg eccentricity, and pRF size to 2/3 of the maximum eccentricity (i.e., 6.67deg half-width radius). Voxels for which the pRF model explained less than 10% of the variance were also excluded from further analyses. Note that due to the absence of stimulation blanks in our retinotopic mapping sequence, we restricted pRF estimation to early visual areas (i.e., V1), where pRF are not too foveally-biased and have relatively small pRF size (Dumoulin and Wandell, 2008). Using this method, we derived reliable retinotopic and pRF size maps, which allowed us to estimate coverage maps of the visual field from V1 voxels (**Fig.6**). Each voxel covers a specific region of the visual field, and many points in the visual field are covered by at least one pRF. First, we combined the pRF center and size estimates from each voxel using 2D Gaussians with peak amplitude normalized to 1. pRF covering regions of the blind field corresponded to pRFs with individual normalized amplitude over blind-field regions higher than 66% (**Fig.6a**). To assess changes in pRF size as a function of pRF eccentricity (**Fig.6f**), the median pRF size was computed for each participant at 4 different eccentricities (2-8deg eccentricity; using ±1deg bins). Then, visual coverage maps were created by representing the highest pRF value at each point of the visual field (Amano et al., 2009; Papanikolaou et al., 2014; Wandell and Winawer, 2015). Given that the peak value of the 2D Gaussian was normalized to 1, the range of values at each point of the subject’s coverage map was between 0 (no coverage) and 1 (full coverage).

### Statistical analyses

Standard parametric tests (i.e., repeated-measures ANOVAs, paired t-tests) were used to assess reliable within-subject differences. In all cases in which the Mauchly’s test of sphericity indicated a violation of the sphericity assumption, Greenhouse-Geisser corrected values were used. Partial eta-squared 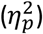 and Cohen’s *d* values were calculated to assess effect size for ANOVAs and paired-sample t-tests, respectively. Multiple linear regressions were used to model the relationship between explanatory variables (e.g., pre-training V1 response and distance from blind-field border) and the dependent outcome (HVF recovery). The coefficient of determination (R^2^) was reported as a measure of the proportion of variance in the dependent variable that is explained by all independent variables, along with the corresponding F-statistic against the constant model. Multicollinearity was assessed by measuring the Variance Inflation Factors (VIF), which indicated no serious correlation between independent variables tested (i.e., VIF between 1.06 and 4.1). Linear regression indicated that the the magnitude of HVF recovery (ΔHVF_prepost_) at blind-field locations was equal to 4.37dB + 7.14dB* pre-training V1 coherence - 0.5dB*blind-field border distance, with a significant interaction term between the two predictors (−1.76dB* pre-training V1 coherence*border distance). Similarly, residual pre-training HVF sensitivity was also predictive of the amount of HVF recovery (ΔHVF_prepost_ = 0.90dB + 4.36dB*pre-training V1 coherence + 0.86dB*pre-training HVF sensitivity).

