## Supplemental Figures for "Changes in perilesional V1 underlie training-induced recovery in cortically-blind patients"

### Supplementary Figures

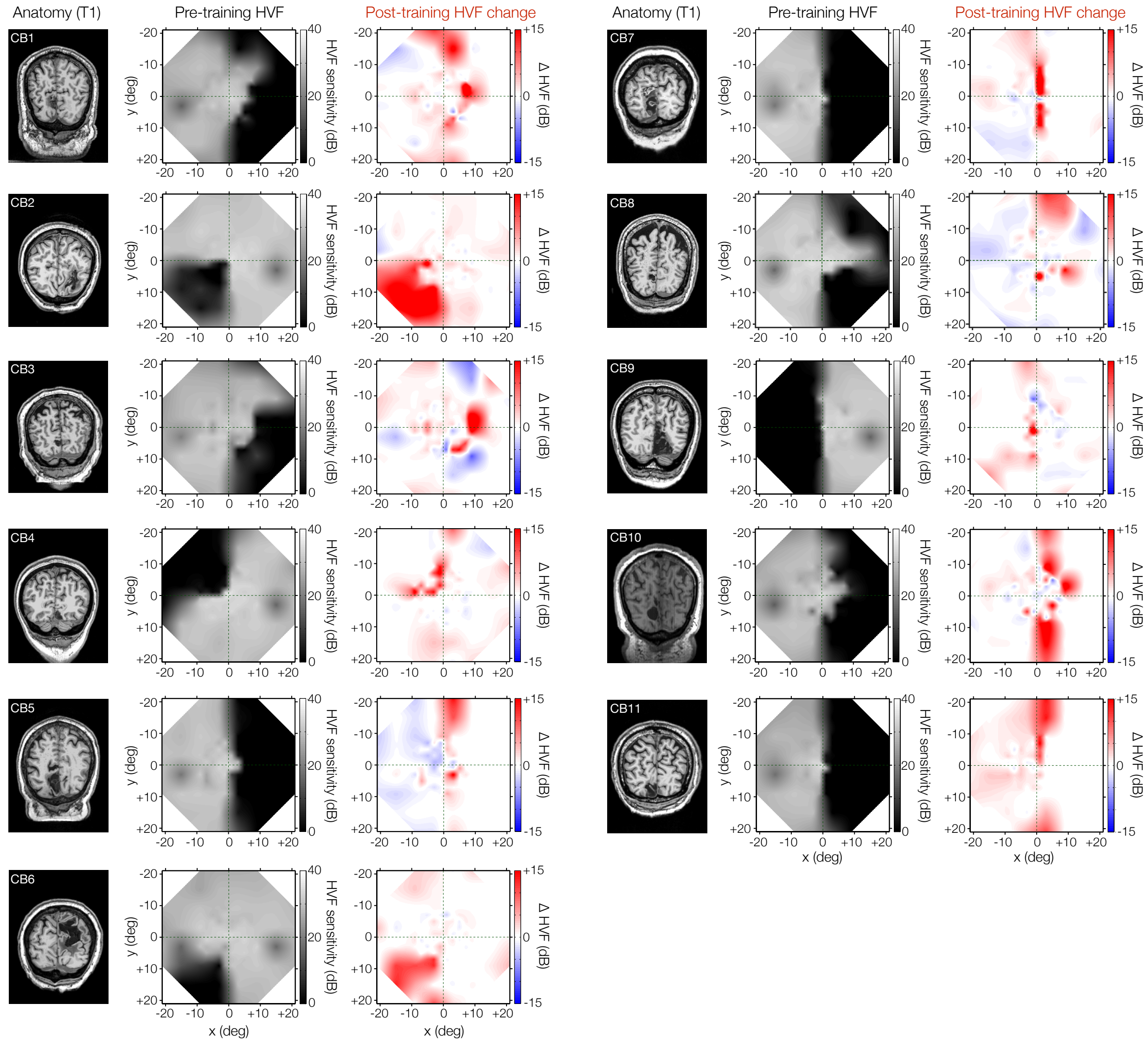


**Figure S1**. *Visual discrimination training improves conscious luminance detection sensitivity in chronic CB patients.* T1-weighted MRI and corresponding baseline (pre-training) composite, binocular Humphrey Visual Fields (HVF; luminance detection sensitivity in dB) for all 11 chronic CB patients over the central ±21 degrees of their HVF field. Prior to training, all CB patients showed homonymous losses of HVF sensitivity (dark regions) within parts of their visual field. Following training, as indicated by HVF change maps, all patients showed improved HVF sensitivity (red), which was greatest within the confines of their original blind-field border. A substantial amount of this recovery usually occured within the inner 11.5 degrees of the visual field (see[16] for full description of training-induced HVF recovery in a larger cohort of chronic CB patients). Areas of sensitivity loss (blue) were also noted in some patients, but only 2 (CB3 and CB9) exhibited significant loss (≥-6dB) in their blind hemifields, along with the larger HVF improvements observed.

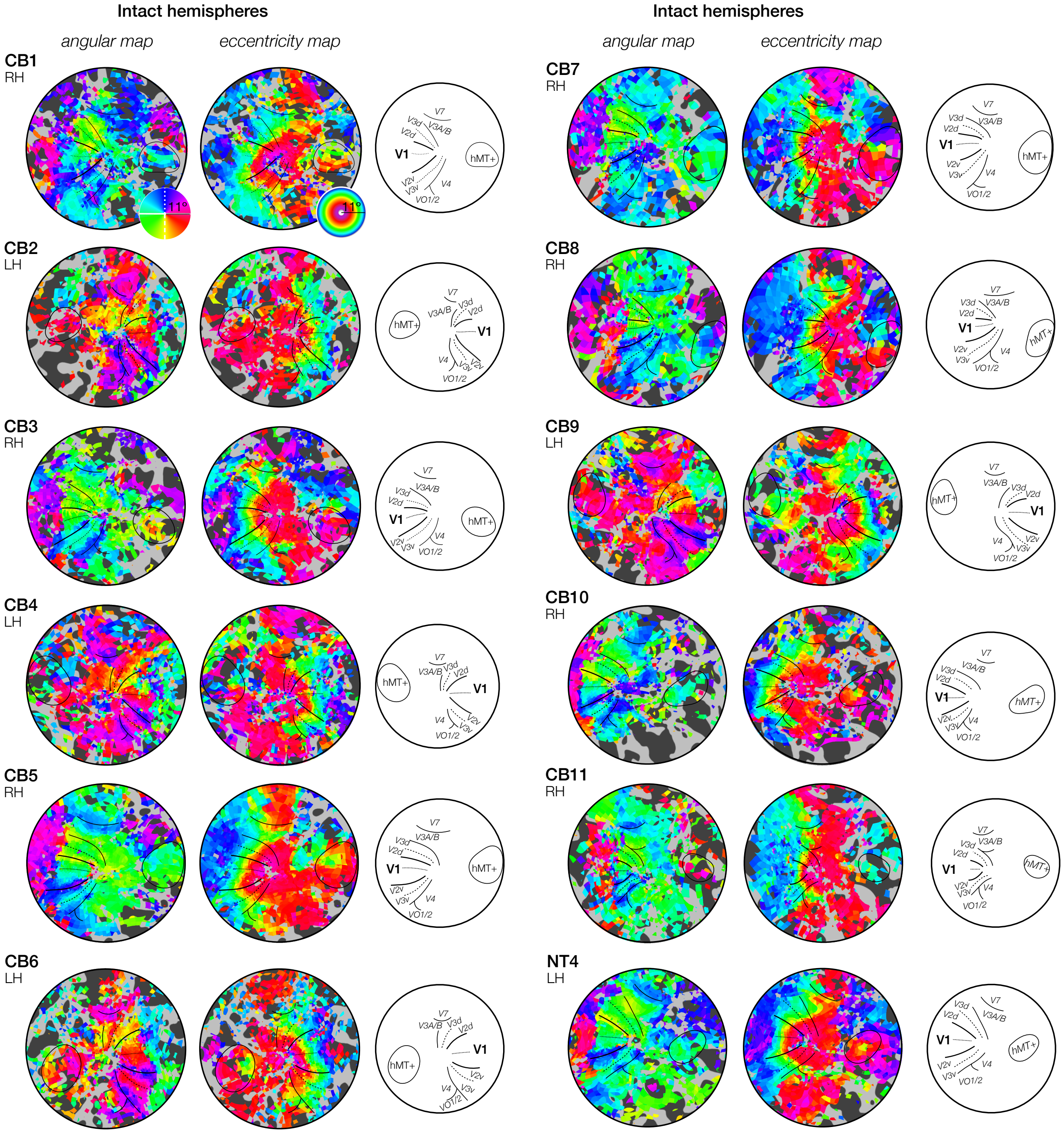


**Figure S2**. *Pre-training retinotopic maps of the intact hemispheres of all 11 chronic CB patients and of one neurotypical (NT) age-matched control.* Visually-evoked activity in the intact hemispheres of all 11 chronic CB patients is retinotopically organized and qualitatively similar to visually-intact controls, both in terms of radial and eccentric representations. The brain hemisphere imaged to obtain these data is indicated for each participant: left hemisphere (LH), right hemisphere (RH).

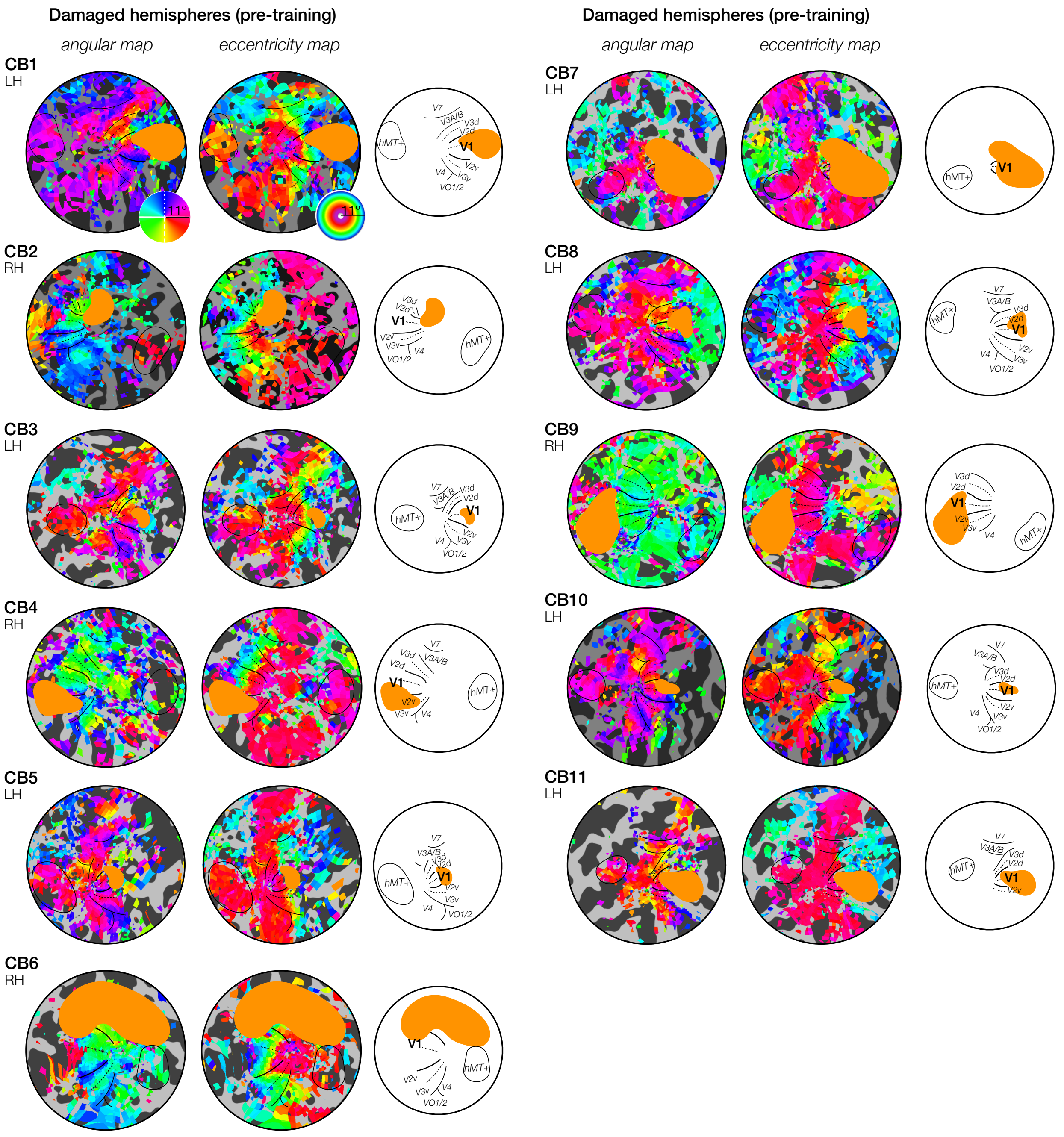


**Figure S3**. *Pre-training retinotopic maps of the damaged hemispheres of all 11 chronic CB patients.* Visually-evoked activity in the damaged hemispheres of chronic CB patients is retinotopically organized, both in terms of radial and eccentric representations, despite the presence of stroke-induced damage (i.e., orange area on maps). The brain hemisphere imaged to obtain these data is indicated for each participant: left hemisphere (LH), right hemisphere (RH).

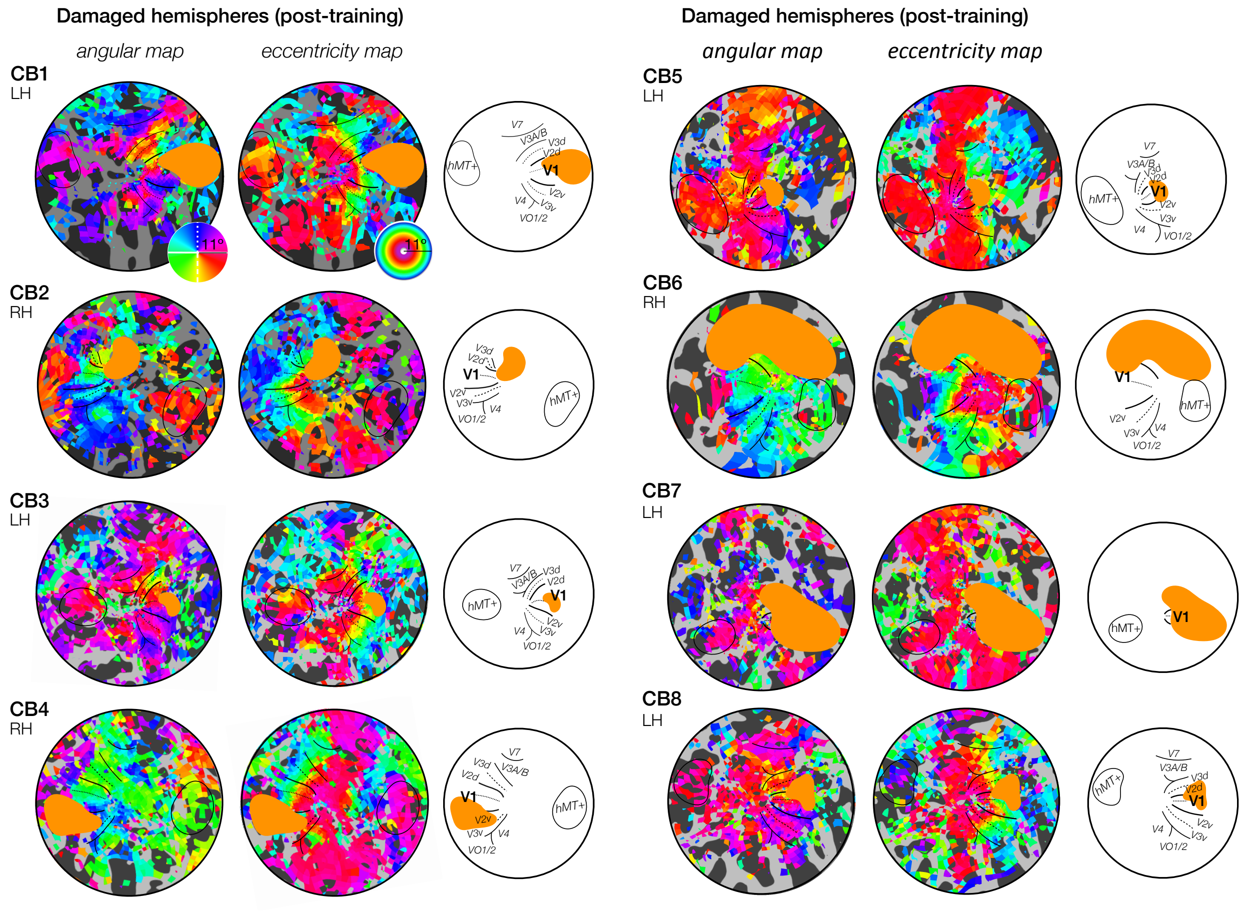


**Figure S4**. *Post-training retinotopic maps of the damaged hemispheres of the 8 chronic CB patients scanned during the post-training visit.* Following training, no coarse change in retinotopic organization was observed. The brain hemisphere imaged to obtain these data is indicated for each participant: left hemisphere (LH), right hemisphere (RH).
